# Resolving the chromatin impact of mosaic variants with targeted Fiber-seq

**DOI:** 10.1101/2024.07.09.602608

**Authors:** Stephanie C. Bohaczuk, Zachary J. Amador, Chang Li, Benjamin J. Mallory, Elliott G. Swanson, Jane Ranchalis, Mitchell R. Vollger, Katherine M. Munson, Tom Walsh, Morgan O. Hamm, Yizi Mao, Andre Lieber, Andrew B. Stergachis

## Abstract

Accurately quantifying the functional consequences of non-coding mosaic variants requires the pairing of DNA sequence with both accessible and closed chromatin architectures along individual DNA molecules—a pairing that cannot be achieved using traditional fragmentation-based chromatin assays. We demonstrate that targeted single-molecule chromatin fiber sequencing (Fiber-seq) achieves this, permitting single-molecule, long-read genomic and epigenomic profiling across targeted >100 kilobase loci with ∼10-fold enrichment over untargeted sequencing. Targeted Fiber-seq reveals that pathogenic expansions of the *DMPK* CTG repeat that underlie Myotonic Dystrophy 1 are characterized by somatic instability and disruption of multiple nearby regulatory elements, both of which are repeat length-dependent. Furthermore, we reveal that therapeutic adenine base editing of the segmentally duplicated γ-globin (*HBG1*/*HBG2*) promoters in primary human hematopoietic cells induced towards an erythroblast lineage increases the accessibility of the *HBG1* promoter as well as neighboring regulatory elements. Overall, we find that these non-protein coding mosaic variants can have complex impacts on chromatin architectures, including extending beyond the regulatory element harboring the variant.

## Introduction

Mosaic variants play a central role in the pathogenesis of Mendelian conditions, cancer, autoinflammatory diseases, and aging via altering the amino acid sequence of protein-coding genes or the gene regulatory elements critical for the appropriate expression of these genes (Poduri et al. 2013; Bamford et al. 2004; Luks et al. 2015; Holzelova et al. 2004; Alriyami and Polychronakos 2021). Although robust tools exist for detecting mosaic variants (Kim et al. 2018; Krishnamachari et al. 2022; Benjamin et al. 2019), quantifying the functional impact of non-coding mosaic variants on overlying chromatin architectures has proven to be more challenging. For example, germline variants that alter chromatin architectures are often detected by identifying ATAC-seq or ChIP-seq peaks with allelically imbalanced read counts. However, by definition, mosaic variants have allelically imbalanced read counts, confounding the ability to accurately disentangle the relative contribution of the variant on chromatin accessibility, especially since the variant allele fraction of a mosaic variant can shift depending on which section of a tissue is profiled. Furthermore, these short-read methods are ill-suited for identifying neighboring regulatory elements that may be altered by a variant as the reads are often less than 150 bp in length, resulting in a myopic view of how mosaic variants impact gene regulatory architectures.

Long-read sequencing allows for the precise identification of single-nucleotide and structural variants across individual multi-kilobase sequencing reads. Moreover, recent advances in single-molecule chromatin fiber sequencing enable the co-identification of genetic variants and chromatin architectures along individual chromatin fibers (Stergachis et al. 2020; Lee et al. 2020; Abdulhay et al. 2020; Shipony et al. 2020; Altemose et al. 2022; Cheetham et al. 2022). For example, Fiber-seq uses non-specific *N*^6^-adenine methyltransferases (m6A-MTases) to selectively mark regions of chromatin accessibility and protein occupancy along individual DNA molecules via m6A-modified bases (Stergachis et al. 2020). m6A-modified bases along with endogenous CpG methylation are then detected using PCR-free long-read sequencing (Marks et al. 2012; Clark et al. 2012; Murray et al. 2012; Loman et al. 2015; Töpfer and Wenger 2023), enabling the single-molecule detection of genetic and chromatin information with the ability to evaluate these features within challenging genomic regions that are known to play a pivotal role in many human diseases (Cooper et al. 2011). We hypothesized that the single-molecule long-read nature of Fiber-seq would be well-suited for investigating the chromatin impact of mosaic variants and sought to create a targeted version of Fiber-seq that would combine the single-molecule and nucleotide precision aspects of Fiber-seq with targeted high-molecular-weight DNA enrichment (i.e. targeted Fiber-seq) (**Fig. 1a**).

**Figure 1.**
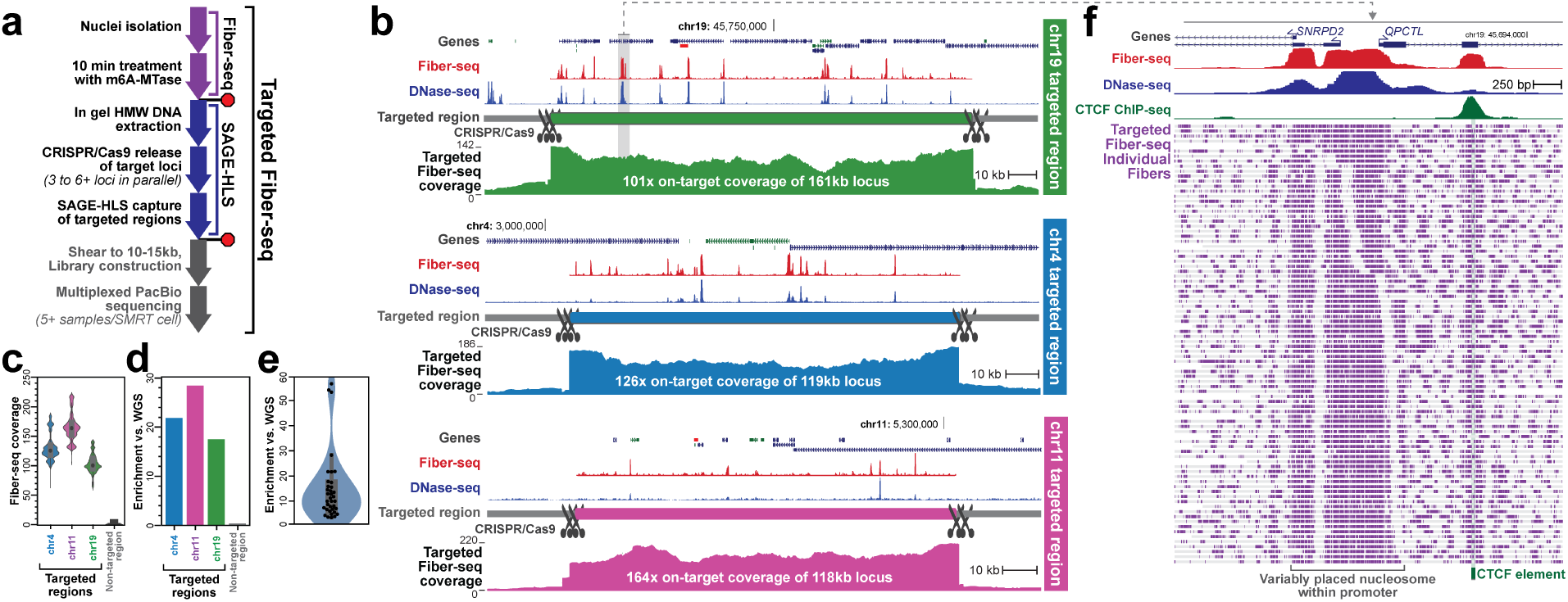
Targeted Fiber-seq methodology. **a)** Schematic of targeted Fiber-seq protocol. **b)** Fiber-seq percent actuation (red), ENCODE DNase I-seq (blue), and per-base targeted fiber-seq coverage are shown across targeted loci. **c)** Violin plot of per-base coverage over the targeted regions, compared to a non-targeted region (chr14:20283833-20402650, hg38). **d)** Relative enrichment of targeted loci relative to WGS (*Methods*). **e)** Relative enrichment across all samples and targets. **f)** Zoom-in of an illustrative locus showing m6A events (purple ticks) across individual fibers.

## Results

To assess the chromatin impact of variants across extended genomic loci, we designed targeted Fiber-seq to simultaneously capture multiple 100-250kb genomic loci using a CRISPR/Cas9 targeting approach. Specifically, four wells of an HLS-SAGE cassette are each loaded with 0.5-1.5 million nuclei treated with m6A-MTase. These samples are then subjected to in-gel DNA extraction and CRISPR/Cas9 release of the targeted loci. The released DNA fragments are then separated and eluted from the gel using pulse-field electrophoresis, sheared to ∼13 kb, barcoded, multiplexed, and then sequenced using single-molecule long-read sequencing (**Fig. 1a**).

As a proof of concept, we first applied targeted Fiber-seq to cultured lymphoblastoid cells (GM04820), which represented 58% of the total mass loaded onto a PacBio Sequel II SMRT cell. A median of 101-164x coverage was achieved across all three targeted loci (**Fig 1b,c**), corresponding to ∼20-fold enrichment compared to whole-genome approaches (**Fig.1d,** *Methods*). A median of 10-fold enrichment was obtained across all samples (**Fig. 1e**).

Comprehensive, deep-coverage epigenetic profiles revealed marked heterogeneity within the chromatin architecture of individual chromatin fibers. For example, although the *SNRPD2/QPCTL* bidirectional promoter was marked by accessible chromatin on each fiber, the boundaries of this accessible promoter element, as well as the positioning of an internal nucleosome varied across the individual reads (**Fig. 1f**), indicating that the precise pattern of accessibility within a promoter can vary quite substantially from fiber to fiber.

Given the potential for targeted Fiber-seq to synchronously resolve alterations in both the genome and chromatin epigenome at single-molecule resolution, we next applied targeted Fiber-seq to evaluate the chromatin impact of the unstable CTG expansion in the 3’ UTR of the gene *DMPK*, which causes Myotonic Dystrophy 1 (DM1) (Brook et al. 1992; Mahadevan et al. 1992; Buxton et al. 1992; Fu et al. 1992; Harley et al. 1992). DM1 is a dominantly inherited trinucleotide repeat disorder characterized by muscle weakness, myotonia, early cataracts, and other symptoms, which become progressively more severe with age. *DMPK* CTG repeat expansions, which are somatically unstable (Wong et al. 1995; Anvret et al. 1993; Ashizawa et al. 1993), disrupt RNA pathways (Ranum and Day 2004; Meola and Cardani 2015; Ravel-Chapuis et al. 2012; Miller et al. 2000; Klesert et al. 1997; Frisch et al. 2001; Hamshere et al. 1997) and locally alter chromatin compaction and CpG methylation (López Castel et al. 2011; Otten and Tapscott 1995; Filippova et al. 2001). However, the extent to which repeat expansions impact chromatin compaction and CpG methylation is not well resolved owing to the inherent limitations of traditional methods. Specifically, the precise boundaries of chromatin accessibility disruptions, the prevalence, and the co-occurrence of these changes on individual chromatin fibers, remain unknown.

To address this, we applied targeted Fiber-seq to enrich for the *DMPK* locus in fibroblasts from four symptomatic individuals in a family with DM1 (**Fig. 2a**). Using reads overlapping the *DMPK* CTG repeats, we quantified repeat instability at single-molecule resolution (**Fig. 2b**, **Supplemental Fig. S1)**. We found that, whereas the affected individual in the first generation of this pedigree had a median pathogenic *DMPK* CTG repeat expansion of 59.5 CTGs and a range of 20 CTG units, *DPMK* CTG repeats in individuals in the 2nd and 3rd generations were consistently >1000 CTGs, with individual chromatin fibers differing by almost 2,000 CTG repeats within an individual donor (**Supplemental Fig. S1)**.

**Figure 2.**
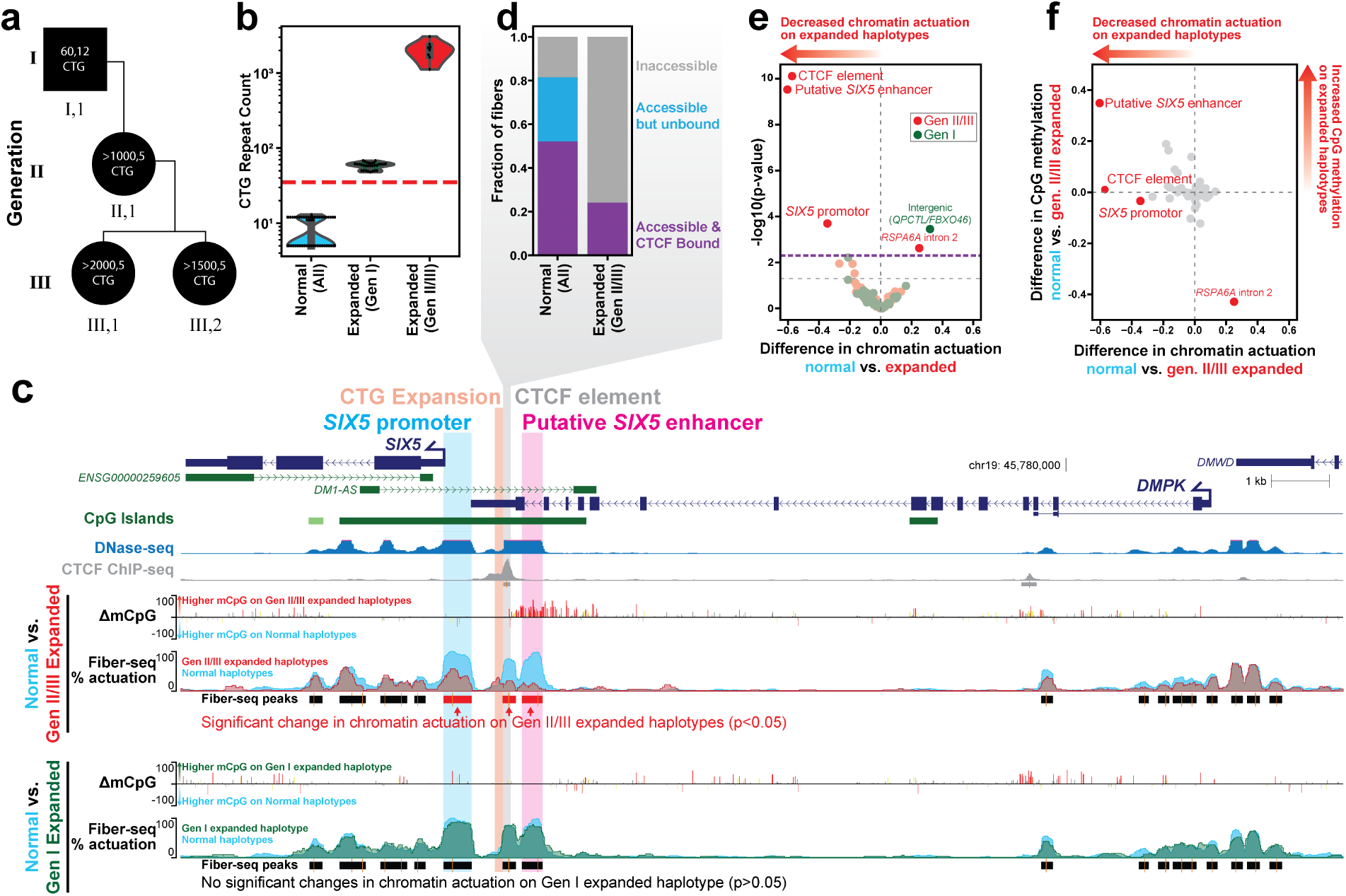
Haplotype-resolved chromatin accessibility and CpG methylation at the *DMPK* 3’ UTR CTG expansion in DM1 fibroblasts. **a)** Family pedigree representing fibroblast donors. Labels inside each shape represent CTG copy number within the pathogenic and normal allele, respectively. **b)** CTG count across sequencing reads that fully span the CTG repeats, grouped by haplotypes. The red dashed line at 35 represents the threshold above which CTG expansions are unstable. **c)** Browser tracks comparing the chromatin architecture of the normal haplotypes to the expanded haplotypes from generations II/III and generation I. The difference in CpG methylation is compared (yellow-p<0.01, orange-p<0.001, red-p<0.0001, Fisher’s exact test), as well as percent actuation of the normal (light blue), generations II/III expanded (red), and generation I expanded (green) haplotypes. Fiber-seq peaks from the normal haplotypes are shown below and are colored to represent a statistically significant decrease in chromatin accessibility on the generation II/III haplotypes (red) compared to normal. There were no significant changes between the normal and generation I (green) expanded haplotype. ENCODE DNase-seq and CTCF ChIP-seq are above in blue and gray, respectively. **d)** CTCF footprinting at the CTG-adjacent CTCF binding site. Footprints were classified as accessible if they were fully overlapped by a methylation sensitive patch, and accessible footprints were classified as CTCF-bound if they did not contain any m6A. (p=0.010, Fisher’s exact test comparing CTCF-bound to unbound [inaccessible plus accessible but unbound]). **e)** Volcano plot of chromatin actuation difference at each accessible peak from the normal haplotype peak set within the targeted locus, compared between normal and expanded fibers from generation I (green) and generations II/III (red). The two peaks with increased accessibility in expanded fibers (upper right quadrant) are likely explained by SNPs local to each region (*Supplemental Note*). P-values were calculated by Fisher’s exact test. The gray dashed horizontal line indicates the nominal significance threshold (p<0.05), and the purple dashed line indicates the Benjamini-Hochberg FDR-corrected significance threshold. **f)** Plot of CpG methylation difference vs. chromatin actuation difference at the peaks described in e. Significant and non-significant points from e are shown in red and gray, respectively.

We next leveraged the underlying highly accurate sequencing information to haplotype-phase reads from each donor within the 160 kb targeted region, thereby identifying whether it arose from chromatin fibers containing the normal or pathogenic repeat expansion (**Fig. 2c**, **Supplemental Fig. S2**, *Methods*). Pathogenic *DMPK* CTG repeat expansions have previously been associated with increased CpG methylation upstream of the CTG repeat, which has been proposed to directly inhibit CTCF occupancy at an upstream element (Filippova et al. 2001). Additionally, decreased DNaseI hypersensitivity of the downstream *SIX5* promoter has also been observed, along with reduced *SIX5* mRNA levels (Klesert et al. 1997). The CpG methylation data obtained via targeted Fiber-seq demonstrates that pathogenic *DMPK* CTG repeat expansions from individuals in the 2nd and 3rd generation are associated with a focal 1 kb hyper-CpG methylated domain upstream of the CTG repeat that extends to the end of the overlapping CpG island. Notably, this hyper-CpG methylated domain does not include the upstream CTCF element, but rather initiates immediately after it (**Fig. 2c**). In contrast, CpG methylation downstream of the *DMPK* CTG repeat is unchanged - indicative of a focal, directional impact of the pathogenic repeat expansion on CpG methylation.

Along the non-pathogenic haplotype, the *DMPK* CTG repeat is bookended downstream by the accessible *SIX5* promoter, and upstream by both a CTCF-bound accessible element and an adjacent accessible element positive for H3K4me1/H3K27ac in ENCODE fibroblast datasets (**Fig. 2c**, **Supplemental Fig. S2**). Of note, this upstream accessible element maps within a region that was previously reported to harbor a putative *SIX5* enhancer, but the exact position of the enhancer within this region had not been determined (Yanovsky-Dagan et al. 2015). Leveraging the paired Fiber-seq chromatin architectures along each fiber, we observed that the putative *SIX5* enhancer is encompassed by the 1kb hyper CpG-methylated domain, and chromatin accessibility is largely ablated across the repeat expansion haplotypes from the 2nd and 3rd generation (**Fig. 2c,e,f**, p-value=7.3e-09, Fisher’s exact test with Benjamini-Hochberg FDR correction). This result demonstrates a direct link between *DMPK* repeat expansion, focal CpG hyper-methylation, and reduced chromatin accessibility at this upstream element.

Although the hyper-CpG methylated domain did not overlap the CTCF binding motif, there was a substantial reduction in chromatin accessibility of the underlying regulatory element in expanded fibers from the 2nd and 3rd generations (p-value=3.8e-09, Fisher’s exact test with Benjamini-Hochberg FDR correction). Utilizing the near single-base pair resolution of targeted Fiber-seq, we classified fibers at the CTCF motif as accessible and CTCF bound, accessible and unbound, or inaccessible (**Fig. 2d)**. Compared to fibers from normal haplotypes which were 52% bound by CTCF, 24% expanded fibers from the 2nd and 3rd generation were CTCF-bound (p-value=0.010, Fisher’s exact test). This reduction in binding was correlated with an increase in nucleosome occupancy overlapping the CTCF binding motif. Together, these results indicate that hyper-CpG methylation of the upstream CTCF element is not required to inhibit CTCF binding on CTG-expanded fibers, as has previously been suggested (Filippova et al. 2001).

We next sought to detect any long-range chromatin impacts associated with these altered elements across the 160kb targeted domain. We observed that of the 96 annotated TSSs within this targeted region, only actuation of the *SIX5* promoter was significantly reduced along expanded fibers from the 2nd and 3rd generations (**Supplemental Fig. S3-4**, p-value=0.028, Fisher’s exact test with Benjamini-Hochberg FDR correction). The canonical *DPMK* TSS had unchanged chromatin accessibility, consistent with findings that suggest post-transcriptional mechanisms drive reduced *DMPK* mRNA levels in DM1 (Hamshere et al. 1997; Frisch et al. 2001). Notably, CpG methylation was unchanged at the *SIX5* promoter, consistent with its altered promoter accessibility being predominantly driven by a loss of an enhancer, as opposed to CpG methylation mediated silencing via the repeat itself (**Fig. 2f**). Together, these findings indicate that pathogenic *DMPK* repeat expansions disrupt chromatin via a focal alteration in upstream CpG methylation and chromatin accessibility, and implicate this upstream enhancer-like element as an enhancer for *SIX5*.

In contrast, we observed that the aforementioned differences in CpG methylation, chromatin accessibility, and CTCF occupancy were not observed along chromatin fibers from the pathogenic allele in the 1st generation, which contains only ∼60 CTG repeats. This suggests that the observed chromatin phenotype of DM1 requires a critical number of CTG repeats, which may be reached either by age-related somatic expansion in adult-onset DM1 or congenitally present in individuals born with large germline CTG repeats.

Next, we sought to apply targeted Fiber-seq to resolve how therapeutic base editing of the fetal γ-globin promoters (encoded by *HBG1/HBG2)* in human CD34^+^-derived erythroid cells affects chromatin structure. Reactivation of fetal γ-globin expression is a promising therapy for β-hemoglobinopathies, as γ-globin can disrupt the formation of HbS polymerization in sickle cell disease and supplement β-globin insufficiency in β-thalassemia (Eaton and Hofrichter 1987; Akinsheye et al. 2011; Bollekens and Forget 1991; Steinberg et al. 2014). The expression of *HBG1* and *HBG2* are repressed in adulthood by the transcriptional regulator BCL11A (Sankaran et al. 2008), and base editing therapies that ablate BCL11A occupancy of the *HBG1* and *HBG2* promoters are in clinical investigation (Beam Therapeutics Inc. 2022). However, our mechanistic understanding of how base editing the *HBG1* and *HBG2* promoters impacts fetal γ-globin expression is duly challenged as base editing is often incomplete, which results in a mixture of edited and unedited haplotypes within a population of cells, and the *HBG1* and *HBG2* genes are situated within a highly similar segmental duplication with 100% sequence identity over the proximal promoter, limiting short-read chromatin methods from uniquely profiling this region (**Supplementary Fig. S5**). To address this, we treated CD34^+^ human hematopoietic stem cells (HSCs) isolated from two healthy donors with a targeted adenine base editor (ABE) to convert A>G within a binding site for BCL11A located -113 bp upstream of both the *HBG1* and *HBG2* TSSs. This same variant is known to cause hereditary persistence of fetal hemoglobin (HPFH) when present in the germline (Amato et al. 2014). We induced differentiation towards an erythroblast lineage and subjected these cells to targeted Fiber-seq, targeting a 118 kb region spanning from the β-globin locus control region (LCR) to the 3’HS1 (**Fig. 3a)**. Notably, *de novo* genome assembly of the Fiber-seq reads from each individual revealed that one of the donors harbored a rare γ-globin gene triplication along one of their haplotypes (**Supplemental Fig. S6-7**). Given the possible confounding features of this triplication event, we removed reads arising from this triplicated haplotype from our downstream analyses.

**Figure 3.**
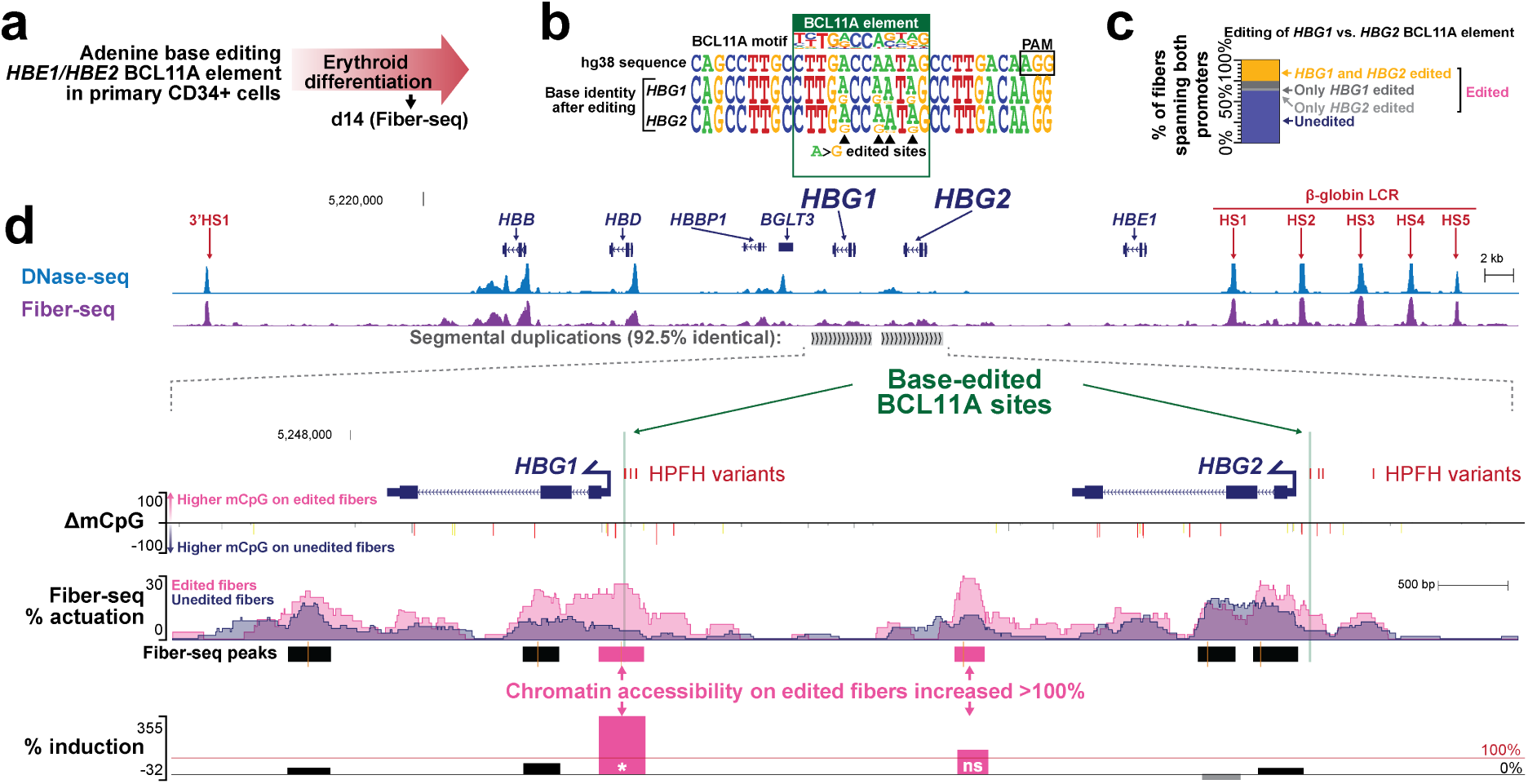
Therapeutic base editing of a BCL11A element within the *HBG1*/*HBG2* promoters. **a)** Schematic of the experimental paradigm. **b)** Sequence logo showing base editing within the BCL11A binding site in the *HBG1* and *HBG2* promoters. Letter height corresponds to the relative base frequency. **c)** Percent of fibers with edited BCL11A sites in both *HBG1* and *HBG2* (yellow), *HBG1* only (dark gray), *HBG2* only (light gray), and neither (blue). The top three categories are grouped as “edited” reads in d. **d)** UCSC browser tracks of ABE-edited CD34^+^-derived erythroid cells. (Top) Comparison of ENCODE DNase-seq of treated multipotent progenitor cells (blue) and chromatin accessibility (Fiber-seq FIRE) of CD34^+^ derived erythroids (purple) across all fibers mapping within the targeted region. (Bottom) Zoom-in with comparison of edited and unedited fibers across *HBG1* and *HBG2*. The difference in CpG methylation is compared (unedited minus edited, yellow-p<0.01, orange-p<0.001, red-p<0.0001, Fisher’s exact test), as well as percent actuation of unedited fibers (blue), and *HBG1* and/or *HBG2* edited fibers (pink). Peak calls for edited reads and percent induction at each peak ([edited percent actuated - unedited percent actuated] / unedited percent actuated) are displayed at the bottom. Pink peaks and bars represent >100% induction. * indicates statistical significance (Fisher’s exact test with Benjamini-Hochberg corrected FDR < 5%, **Supplemental Fig. S9**).

As a first step, we evaluated single-molecule editing efficiency across the β-globin locus, as well as for altered chromatin architectures induced along the base edited chromatin fibers (**Fig. 3a**). We observed that in addition to the targeted -113 A>G edit, edited fibers almost exclusively contained additional A>G edits in the immediate vicinity (**Fig. 3b**). Although only 34% and 32% of fibers demonstrated base editing at the *HBG1* and *HBG2* promoters, respectively (*Methods*), we found that if one of the promoters was edited, the other promoter on that same fiber was also significantly likely to be edited (p < 0.00001, Fisher’s exact test), suggesting that editing efficiency is largely determined by transduction and expression of the ABE (**Fig. 3c**). As such, we grouped fibers edited at *HBG1* and/or *HBG2* fibers for further analyses.

Importantly, the long-reads obtained by Fiber-seq enable the unique mapping of CpG methylation, chromatin architectures, and ABE-induced base changes to each of the segmentally duplicated *HBG1* and *HBG2* genes (**Fig. 3d**, **Supplemental Fig. S8**). Comparison of the CpG methylation patterns between the edited and unedited reads demonstrated that base editing of the *HBG1* and *HBG2* promoters resulted in a significant reduction in CpG methylation over the entire span of both the *HBG1* and *HBG2* genes.

In addition to this broad change in CpG methylation, we also saw focal changes in chromatin accessibility. Specifically, base editing of the *HBG1* and *HBG2* promoters resulted in a >100% increase in chromatin accessibility of the *HBG1* promoter (Fisher’s exact test p-value=0.00030, p-value=0.0033 after Benjamini-Hochberg FDR correction), as well as another element located 2.5 kb upstream of the *HBG1* promoter (Fisher’s exact test p-value=0.024, p-value=0.13 after Benjamini-Hochberg FDR correction) (**Fig.3d, Supplemental Fig. S9)**. In contrast, chromatin accessibility at the four other accessible elements within the *HBG1/HBG2* segmental duplication exhibited non-significant changes along the edited fibers, including the *HBG2* promoter, which demonstrated only a 36% increase in accessibility along the edited fibers. Notably, the four non-promoter regulatory elements that we identified within the *HBG1/HBG2* segmental duplication were enriched for histone modifications associated with active regulatory elements in hematopoietic cells, and chromatin accessibility of these elements was coupled with the accessibility of the *HBG1/HBG2* promoters along single molecules (**Supplemental Fig. S8**), suggesting a putative role as *HBG1/HBG2* enhancer elements. Together, these findings demonstrate that base editing of the *HBG1* and *HBG2* -113 A>G sites preferentially occurs along the same chromatin fiber and induces chromatin accessibility at *HBG1,* likely in cooperation with adjacent enhancer-like elements.

## Discussion

Overall, we demonstrate that targeted Fiber-seq enables the production of targeted long-read sequencing chromatin maps to resolve the heterogeneity of genetic and chromatin architectures with single-molecule precision. Importantly, as targeted Fiber-seq enables the synchronous measurement of DNA sequence, CpG methylation, and chromatin accessibility along the same +10kb molecule of DNA, we can directly disentangle the functional impact of heterogeneously present genetic variants on neighboring gene regulatory programs. Application of this to the *DMPK* and *HBG1/HBG2* loci demonstrated that precise genetic changes can result in complex alterations in chromatin accessibility that extend beyond the genetically modified element or the region with disrupted CpG methylation - alterations that would be largely hidden from short-read allelic imbalance or long-read CpG methylation methods.

With combined and phased chromatin accessibility and CpG methylation on the same chromatin fibers, our results provide new insight into the chromatin phenotype of DM1. First, our results substantiate the seemingly contradictory findings that chromatin accessibility of the *SIX5* promoter is decreased (Klesert et al. 1997; Otten and Tapscott 1995) while CpG methylation is unchanged (López Castel et al. 2011). More broadly, this result highlights that although CpG methylation and chromatin accessibility are correlated (ENCODE Project Consortium 2012), they are not interchangeable. Second, we show for the first time that reduced CTCF binding to the element located directly upstream of the CTG repeat can occur without a corresponding increase in overlapping CpG methylation. *In vitro* binding studies previously demonstrated that CpG methylation of the CTCF binding motif inhibited CTCF binding (Filippova et al. 2001), and so it was proposed that CpG methylation is the mechanism by which CTCF binding is reduced *in vivo*. In support of our findings, more recent studies using Oxford Nanopore (ONT) sequencing also show variable CpG methylation of the CTCF binding element in individuals with DM1 (Rasmussen et al. 2022). However, these ONT studies are incapable of co-measuring CTCF occupancy. Co-measurements of CpG methylation, chromatin accessibility, and single-molecule CTCF footprinting enabled us to eliminate any confounding factors and directly disentangle this result.

In addition, we demonstrate the utility of targeted Fiber-seq for resolving both genetic and gene regulatory alterations within complex genomic regions such as segmental duplications. Specifically, we were able to precisely disentangle the genetic and chromatin architectures of the *HBG1* and *HBG2* genes, demonstrating that the γ-globin -113 A>G variant that is known to cause HPFH results in the focal increase in chromatin accessibility at the γ-globin gene promoters, as well as at adjacent regulatory elements. Furthermore, we were able to perform *de novo* genome assembly on our data to show that one of the donors harbored a rare γ-globin gene triplication along one of their haplotypes. This later observation enabled us to selectively remove reads arising from this triplicated haplotype from our analyses above, reducing a possible confounding feature that would have been largely hidden from short-read-based chromatin methods.

Our results provide mechanistic insights into the pathogenesis of both DM1 and HPFH, precisely delineating enhancer elements that potentially play a critical role in both of these conditions. Specifically, we identify that pathogenic *DMPK* repeat expansions disrupt an upstream accessible chromatin element with enhancer-like activity, and implicate this upstream enhancer-like element as an enhancer for *SIX5*, providing a mechanistic basis for altered *SIX5* expression, which is one of the hallmark features of DM1. We also found that the size of the CTG expansion was correlated with both somatic instability and disruptions in chromatin architecture, raising the possibility that a common mechanism might contribute to both processes. Notably, these findings were performed in patient-derived fibroblasts, and further studies using primary tissues from patients will be helpful to further evaluate this mechanism.

Together, these findings emphasize the benefit of targeted single-molecule chromatin accessibility assays, such as targeted Fiber-seq, to fully capture the genetic and functional impact of germline and mosaic variants. We anticipate that further advances in target capture enrichment will build upon the benefits delineated in this manuscript. Furthermore, by unraveling the chromatin basis of these two disease-associated variants, this study highlights additional potential therapeutic epigenetic targets for both DM1 and β-hemoglobinopathies.

## Methods

### Cell Line Culture

The following cell lines/DNA samples were obtained from the NIGMS Human Genetic Cell Repository at the Coriell Institute for Medical Research: GM04820 (lymphoblastoid), GM06076 (fibroblast), GM04601 (fibroblast), GM04602 (fibroblast), GM04608 (fibroblast). Lymphoblastoid cells were maintained in suspension in IMDM media supplemented with 10% FBS (HyClone, cat. no.SH30396.03IH25-40) and antibiotic (100 I.U/mL penicillin, 100ug/mL streptomycin, Gibco, cat. no. 15140122) at 37°C and 5% CO2 in T-75 flasks. Cells were split 1:10 every 3-4 days. Fibroblasts were maintained in DMEM media with the same supplementation and incubation as above. Cells were split 1:4 every 5-7 days with 0.25% trypsin-EDTA (Gibco, cat. no. 25200056).

### CD34^+^ cell culture and transduction

CD34^+^ cells from G-CSF-mobilized healthy adult donors were provided by the Fred Hutch Cell Processing Facility. The cells were recovered from frozen stocks and incubated overnight in StemSpan H3000 medium (STEMCELL Technologies, Vancouver, Canada) supplemented with penicillin/streptomycin, Flt3 ligand (Flt3L, 25 ng/mL), interleukin 3 (10 ng/mL), thrombopoietin (TPO) (2 ng/mL), and stem cell factor (SCF) (25 ng/mL). Cytokines and growth factors were from Peprotech (Rocky Hill, NJ). CD34^+^ cells were transduced in low-attachment 6-well plates with an all-in-one base editing vector HDAd-ABE8e-sgHBG2 targeting the BCL11A-binding sites in the *HBG1*/*2* promoters for fetal hemoglobin reactivation (Li et al. 2022).

### *In vitro* erythroid differentiation of CD34^+^ cells with O^6^BG/BCNU selection

Differentiation of human CD34^+^ cells into erythroid cells was done based on the protocol developed by Douay et al. (Douay and Giarratana 2009). In brief, in step 1, cells at a density of 10^4^ cells/mL were incubated for 7 days in IMDM supplemented with 5% human plasma, 2 IU/mL heparin, 10 g/mL insulin, 330 g/mL transferrin, 1 M hydrocortisone, 100 ng/mL SCF, 5 ng/mL IL-3, 3 U/mL erythropoietin (Epo), glutamine, and penicillin/streptomycin. In step 2, cells at a density of 1×10^5^ cells/mL were incubated for 3 days in IMDM supplemented with 5% human plasma, 2 IU/mL heparin, 10 g/mL insulin, 330 g/mL transferrin, 100 ng/mL SCF, 3 U/mL Epo, glutamine, and Pen/Strep. In step 3, cells at a density of 1×10^6^ cells/mL cells were incubated for 4 days in IMDM supplemented with 5% human plasma, 2 IU/mL heparin, 10 g/mL insulin, 330 g/mL transferrin, 3 U/mL Epo, glutamine, and Pen/Strep. For the enrichment of transduced cells, 48 hours post-transduction, CD34^+^ cells were incubated with 50 M O⁶-Benzylguanine (O^6^BG) for one hour. Without washing, 35 M Carmustine (BCNU) was added for 2.5 more hours incubation, after which cells were washed and resuspended in fresh medium. Both drugs were purchased from Millipore/Sigma and freshly prepared. O^6^BG/BCNU selection was used to enrich for edited cells in samples PS00208.erythroid1, PS00209.erythroid2, and PS00316.erythroid1.

### Fiber-seq

In-house Hia5 preparation and fiber-seq were performed as described (Stergachis et al. 2020). Briefly, 2-4×10^6^ cells were washed with PBS, lysed in lysis buffer (15mM Tris-Cl, pH 8, 15mM NaCl, 60 mM KCl, 1mM EDTA pH 8, 0.5mM EGTA pH 8, 0.5 mM spermadine, 0.025% IGEPAL) and treated with Hia5 at 100U per million cells (25°C, 10 min). The reaction was split into four equal aliquots, and SDS was added to a final concentration of 2%. The volume of each aliquot was adjusted to 70 μL with suspension buffer M2 (Sage Science). Samples used in this study are described in **Supplemental Table S1**.

### crRNA Design

Three crRNAs per side per target were designed within a ∼3 kb span with attention to avoid annotated repeat elements or common variants. The following target windows represent the region flanked by the innermost crRNAs: chr4:3006837-3125654 (chr4 target), chr11:5186600-5304585 (chr11 target), and chr19:45664901-45825746 (chr 19 target). crRNAs were synthesized by IDT. crRNA sequences are detailed in **Supplemental Table S2**.

### HLS-CATCH

HLS-CATCH was performed according to the manufacturer’s protocol (Sage Science), using the Non-Core Workflow “CATCH 100-300Kb extr3h enhINJ Sep3h”, with modifications as described: For the extraction step, post fiber-seq nuclei aliquots (∼6×10^5^-1×10^6^ cells/lane) were added to HLS-Sage sample wells, and 210 μL of HLS lysis buffer A was added to the reagent wells. Cassettes were sealed and run for 3h, 55 V with wave index 1-1. Meanwhile, crRNAs (3 per target per side) were diluted to 100 μM in nuclease-free duplex buffer (IDT cat#11-01-03-01) and annealed and prepared as specified, using NEB Cas9 (M0386M) or NEB EnGen Spy Cas9 HF1 (M0667M). The Cas9 reaction mix was injected for 2min, 80V with wave index 2-1 and incubated for 30 minutes at RT. The reagent well was replaced with lysis buffer A and ran for 3h, 55V with a wave index 3-2. The sample was eluted for 1.5h, 50V with wave index 3-1.

### Library Preparation and PacBio sequencing

Post-CATCH samples were purified with 0.5x volume of Ampure PB beads and eluted in 52 μL of elution buffer (PacBio 101-633-500). 1 μL was used to quantify sample concentration using a Qubit dsDNA HS Assay according to the manufacturer’s protocol. Samples were sheared to 10-15 kb with two passes (one inversion) through a Covaris gTUBE at 3200rpm for 2-4 minutes per pass in an Eppendorf 5424R centrifuge. ∼50 μL per sample was recovered and barcoded using the SMRTbell® Express Template Prep Kit 2.0(Pacbio) with 5 μL of barcoded adapter from the Barcoded overhang adapter kit 8A/B for Sequel II sequenced samples or the SMRTbell® prep kit 3.0 with the SMRTbell adapter index plate 96A, according to the protocol for each kit with adjustment for larger sample volume. Samples were sequenced by UW PacBio Sequencing Services on a Sequel II SMRT Cell 8M with a 30-hour movie or on a Revio SMRT Cell 25M with a 24-hour movie.

### Fiber-seq Processing

Circular consensus sequence reads were generated from raw PacBio subread files using PacBio CCS (https://ccs.how/) with chunking and average kinetics information included. Chunks were combined with pbmerge (https://pbbam.readthedocs.io/en/latest/tools/pbmerge.html). CCS reads were demultiplexed with lima (https://lima.how/). Demultiplexed bam files were labeled with CpG methylation using Primrose/Jasmine (Töpfer and Wenger 2023), keeping kinetics. m6A and nucleosome calls were generated with fibertools using a 55 base pair nucleosome threshold (Jha et al. 2024). A small percentage of reads with the proportion of methylated adenine to all adenines below 0.02 or above 0.4 were excluded from further analysis. Scripts to filter m6A reads by m6A proportion are available here (https://github.com/StephanieBohaczuk/Targeted-Fiber-seq/tree/main/m6a_percent_filtering).

### Alignment

Reads were aligned to hg38 with pbmm2, which revealed that one of the two CD34^+^ cell donors had a heterozygous duplication of *HBG2* (**Supplemental Fig. S6-7**). To resolve this, a custom assembly of the β-globin locus was created with a separate sample from the same donor (PS00148) using Hifiasm (Cheng et al. 2021) with the -f0 flag to create an assembly from reads aligning to chr11:5,180,424-5,305,559 in hg38. Hifiasm yielded two contigs: A 133 kb contig containing one copy each of *HBG1* and *HBG2* (2γ haplotype) and a 126 kb contig containing one *HBG1* copy and two *HBG2* copies (3γ haplotype). The donor-specific assembly is available here (https://github.com/StephanieBohaczuk/Targeted-Fiber-seq/tree/main/ABE_hifiasm_assembly), and NucFreq plots(Vollger et al. 2019) of alignment to hg38 and the donor-specific assembly are shown in **Supplemental Fig. S7**. Reads from this donor aligning to chr11:5,186,600-5,304,585 in hg38 were remapped to the Hifiasm custom genome and filtered for primary alignments only with MAPQ score > 10. Reads mapping to the 2γ haplotype were filtered from the hg38 alignment and merged with all reads from the second donor, who was homozygous for the 2γ haplotype. Coverage of the 3γ haplotype was insufficient for follow-up analyses.

### Phasing

DM1 reads were phased separately for each sample. First, DeepVariant(Poplin et al. 2018) was used to identify variants and produce a VCF file. Reads were phased with HiPhase(Holt et al. 2024) using the k-mer variant phasing pipeline with default settings (https://github.com/mrvollger/k-mer-variant-phasing).

### Fiber-seq Chromatin Accessibility and peak actuation comparisons

To assess chromatin percent actuation, we used the Fiber-seq Inferred Regulatory Element (FIRE) pipeline version 0.0.4 (https://github.com/fiberseq/fiberseqfire)(Vollger et al. 2024), a method that uses a semi-supervised machine learning algorithm to predict the likelihood that a methylation-sensitive patch is a regulatory element on individual chromatin fibers. The configuration files used to run FIRE are available on GitHub (https://github.com/StephanieBohaczuk/Targeted-Fiber-seq/tree/main/FIRE_config). The fiber-seq chromatin percent actuation tracks in **Fig. 1**, **2**, and 3 represent the percent of fibers with FIRE elements at each base pair. Peaks are called with min_frac_accessible: 0.15. Reads were grouped as follows for FIRE analysis: PS00118 (**Fig. 1**); normal haplotypes from all DM1 samples (PS00150, PS00151, PS00152, PS00153, PS00442, PS00443, PS00444), expanded haplotypes from generation II/III DM1 donors (PS00150, PS00151, PS00152, PS00153, PS00443, PS00444), expanded haplotype from generation I DM1 donor (PS00442) (**Fig. 2**); Edited and unedited 2γ-haplotype from CD34^+^ donor 1 and both haplotypes from CD34^+^ donor 2 (PS00196, PS00208, PS00209, PS00316) (**Fig. 3c**, top track in purple), edited reads from 2γ-haplotype from CD34^+^ donor 1 and both haplotypes from CD34^+^ donor 2 (PS00196, PS00208, PS00209, PS00316), unedited reads from 2γ-haplotype from CD34^+^ donor 1 and both haplotypes from CD34^+^ donor 2 (PS00196, PS00208, PS00209, PS00316) (**Fig. 3**).

Fraction actuation over each peak was computed as the number of reads with a FIRE score <= 0.1 anywhere within the ∼100-200 bp peak divided by the total number of reads mapped within the peak. For DM1 analyses, normal and expanded peaks were compared across peaks called for normal haplotypes from all DM1 samples (**Fig. 3c**). For comparison of edited and unedited CD34^+^-derived erythroids, edited and unedited peaks were compared across all peaks called for the edited haplotype. Fisher’s exact test was performed in python (scipy.stats.fisher_exact) with Benjamini-Hochberg FDR correction (scipy.stats.false_discovery_control) using FDR < 5% for statistical significance. The FDR-corrected significance threshold was calculated as 0.05*(rank of the smallest significant FDR-corrected p-value + 1)/total number of comparisons, corresponding to the p-value that would have been required for the smallest p-value point that did not reach FDR-corrected significance to be significant. In **Fig. 2e** and **Fig. S4**, the FDR-corrected significance threshold is plotted for Generation II/III samples, but Generation I points above and below this line are accurately classified as significant or non-significant, respectively, following FDR correction. Percent induction due to editing (**Fig. 3c**) is calculated as (edited percent actuated - unedited percent actuated) / unedited percent actuated. Scripts to reproduce analyses and figures are available on GitHub (https://github.com/StephanieBohaczuk/Targeted-Fiber-seq/tree/main/compare_perc_actuation).

### CpG methylation

To compare CpG methylation, the haplotype tag (1 or 2) was used to label reads within each bam, and then bam files to compare were merged. ‘aligned_bam_to_cpg_scores’ (Pacific Biosciences 2023) was run with default settings using the “pileup_calling_model.v1.tflite” model. For CpG differences tracks (**Fig. 2c**, **3d**), MethBat (Holt and Saunders 2023) was used to calculate the difference in CpG methylation and an associated p-value at each CpG (i.e. 1 base pair input regions). CpG sites that were present as heterozygous variants (>10% frequency) were excluded. For CpG methylation over peaks (**Fig. 2f**), peaks were used as input regions. Scripts to reproduce CpG tracks are available on GitHub (https://github.com/StephanieBohaczuk/Targeted-Fiber-seq/tree/main/CpG_tracks).

### Coverage and enrichment

Coverage was calculated at each base pair within the targeted loci. Enrichment was calculated as median sequencing depth/(mass fraction × expected WGS coverage), where mass fraction is sample mass/total mass loaded onto the multiplexed SMRT cell, and an expected WGS coverage of 10 and 30 were used for Sequel II and Revio, respectively. Scripts to reproduce analyses are available on GitHub. (https://github.com/StephanieBohaczuk/Targeted-Fiber-seq/tree/main/coverage).

### DM1 repeat expansion length

The CTG/CAG repeat was counted on reads that were anchored both 5’ and 3’ of the repeat, i.e. those where either the primary alignment mapped both 5’ and 3’ of the repeat, or a primary alignment mapped either 5’ or 3’ and a supplementary alignment mapped to the missing side. The length of the expansion was calculated by finding the position of the first and last CAG within the repeat region, subtracting these positions to find the total length of bases, and then dividing by three for the length in CAG units. A custom script to reproduce this analysis is available on GitHub (https://github.com/StephanieBohaczuk/Targeted-Fiber-seq/tree/main/DM1_CAG).

### CTCF footprinting

CTCF footprinting was computed using a custom script, which assigns fibers as “accessible and CTCF bound” if the core of the CTCF binding site (Yin et al. 2017) (chr19:45770329-45770342 in hg38) is fully overlapped by a methylation sensitive patch (MSP) but does not contain any m6A calls within, “accessible but unbound” if the core is fully overlapped by an MSP but does contain m6A calls within, or “inaccessible” if the core is overlapped by a nucleosome. Statistics represent Fisher’s exact test comparing “accessible and CTCF bound” to unbound (i.e. “accessible and unbound” + “inaccessible”). A custom script to reproduce this analysis is available on GitHub (https://github.com/StephanieBohaczuk/Targeted-Fiber-seq/tree/main/DM1_CTCF).

### Identification of ABE edits

Reads mapping to the edited BCL11A site (Li et al. 2022) within the *HBG1* and/or *HBG2* promoters were categorized as “unedited” if they contained no indels or mismatches within a window of -4/+5 bp from the intended edit site located 113 bp upstream of the *HBG1/2* TSS. Reads were categorized as “edited” if they contained the expected A>G (T>C in the reference genome), with other edits within the window also tolerated. Reads not classified as “edited” or “unedited” were excluded from further analyses. Scripts to classify reads by editing status are available on GitHub (https://github.com/StephanieBohaczuk/Targeted-Fiber-seq/tree/main/ABE_editing).

### ENCODE tracks

The following ENCODE (ENCODE Project Consortium 2012) tracks were included: Dnase-seq of GM12878, ENCFF960FMM (**Fig. 1b,f**); DNase-seq of fibroblasts, ENCFF302JEV, CTCF ChIP-seq of fibroblasts, ENCFF080HIA (**Fig. 2c**); DNase-seq of hematopoietic multipotent progenitor cell treated with interleukin-3 for 8 days, kit ligand for 8 days, hydrocortisone succinate for 8 days, erythropoietin for 8 days, ENCFF800PMS (**Fig. 3c**).

### Co-Accessibility and Codependency

Co-accessibility between each pair of peaks was quantified by evaluating fibers that 1) entirely overlapped each of the two peaks and 2) had at least 46 bp between the read ends and peak edges. Fibers were classified as accessible at a peak if they contained a FIRE element (FIRE score <= 0.05) that overlapped the peak by >= 1 bp. Overlaps were computed using Bedtools intersect (v2.31.0) (Quinlan and Hall 2010). Co-accessibility data was filtered to only include peak pairs that were overlapped by a minimum of 20 fibers. Codependency scores were calculated as the difference between the observed and expected co-accessibility given the null hypothesis that the actuation of each peak is independent of the other. The expected co-accessibility was calculated as the product of the actuation proportion at each of the two peaks. The resulting scores were multiplied by 4 to scale the range of possible values from -1 (only one peak is actuated per fiber) to +1 (actuation at both peaks for a given fiber). Scripts to reproduce this analysis is available on GitHub (https://github.com/StephanieBohaczuk/Targeted-Fiber-seq/tree/main/ABE_codependency).

### Data access

Sequencing data generated in this study have been submitted to the NCBI BioProject database (https://www.ncbi.nlm.nih.gov/bioproject/) under accession number PRJNA1125891.

### Code availability

The code to reproduce figures and analyses in this study is available on GitHub https://github.com/StephanieBohaczuk/Targeted-Fiber-seq.

### Competing interest statement

A.B.S. is a co-inventor on a patent relating to the Fiber-seq method (US17/995,058). A.L. is an academic co-founder of Ensoma Inc. The remaining authors declare no competing financial interests.

## Acknowledgments

The authors thank Stephen Tapscott for discussion of the Myotonic Dystrophy 1 data, Shane Neph for assisting in data submission to the NCBI Bioproject database, and Chris Boles for technical guidance on the HLS-CATCH method.

A.B.S. holds a Career Award for Medical Scientists from the Burroughs Wellcome Fund and is a Pew Biomedical Scholar. This study was supported by National Institutes of Health (NIH) grants 1DP5OD029630, and UM1DA058220 to A.B.S., as well as a UW ADRC Developmental Project (NIH grant P30AG066509) to A.B.S.. M.R.V. and S.C.B. were supported by a training grant (T32) from the NIH (2T32GM007454-46).

## Author contributions

Conceptualization and design: S.C.B. and A.B.S. Experimental design and execution: S.C.B., Z.J.A.., C.L., B.J.M., J.R., K.M.M., T.W., Y.M., A.L., and A.B.S. Computational experiments: S.C.B., Z.J.A., E.G.S., M.O.H., and M.R.V. Data generation: S.C.B., Z.J.A., C.L., J.R., K.M.M., and A.B.S. Manuscript writing: S.C.B. and A.B.S., with input from all authors.

## Supplemental Figures

**Figure S1.**
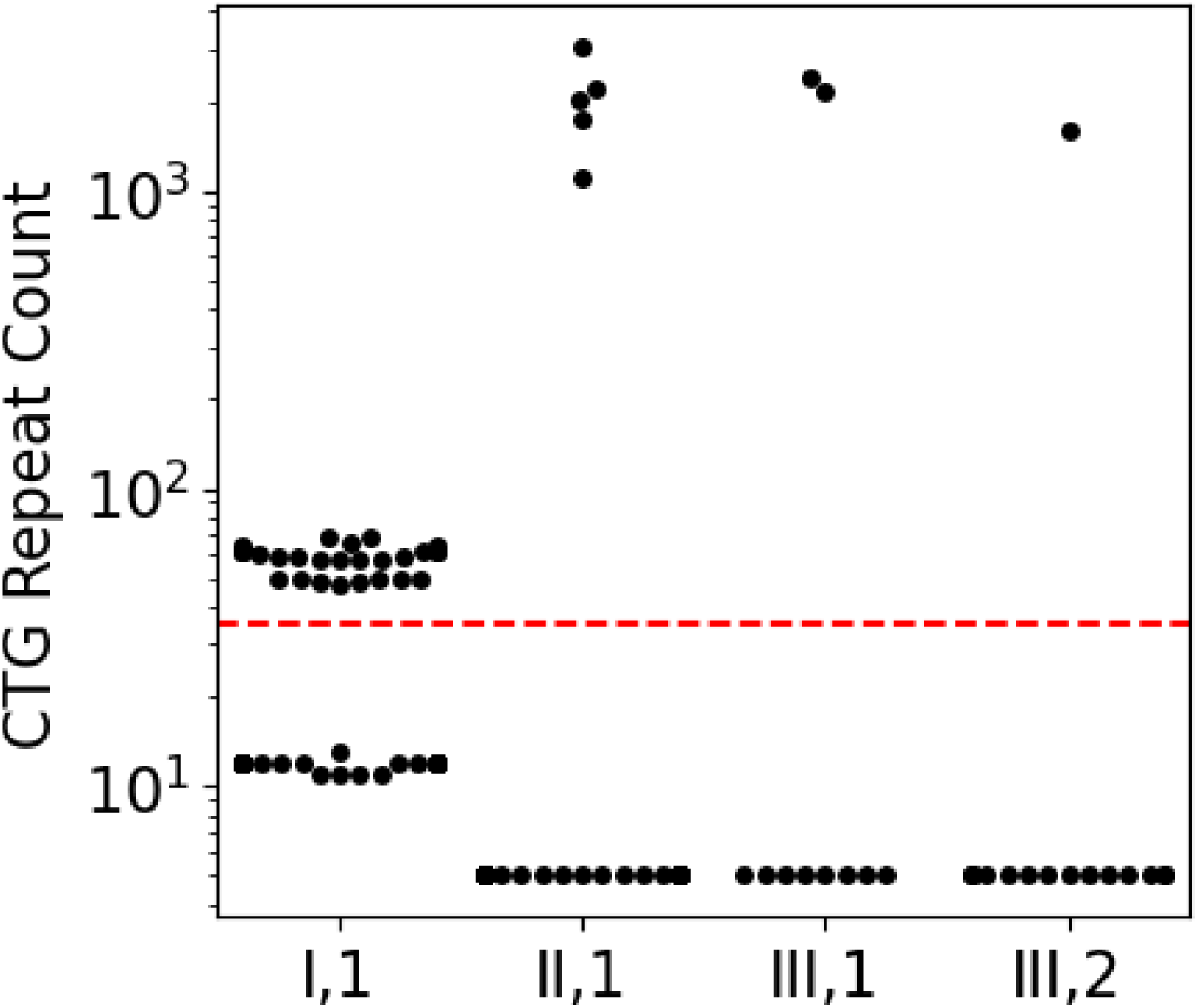
CTG repeat count per donor. Swarm plot of CTG repeat count in reads fully spanning the CTG repeat, separated by individual donor.

**Figure S2.**
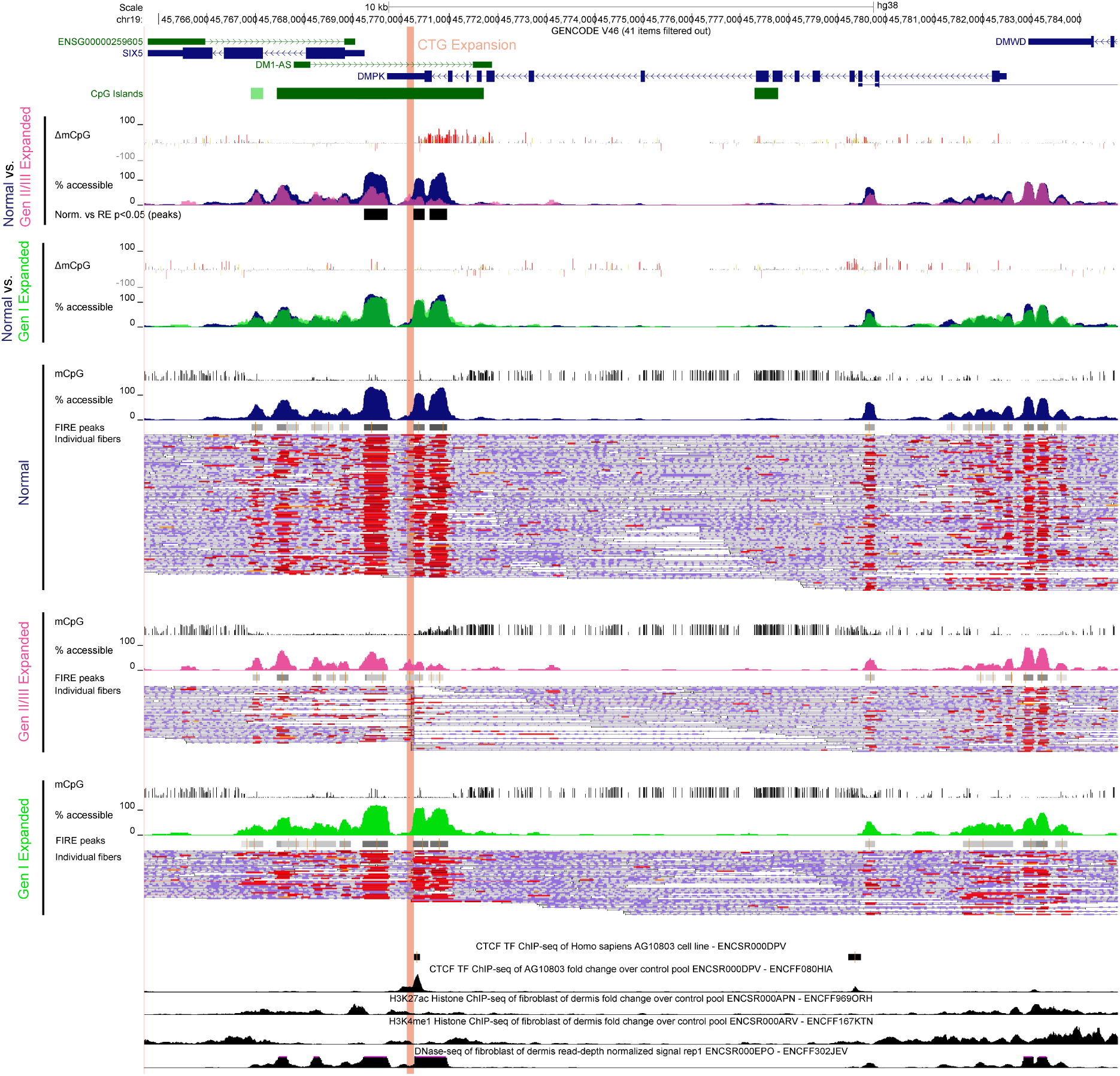
Targeted Fiber-seq of DM1 fibroblasts at *DMPK*. The top two panels show a comparison of mCpG and actuation between normal and expanded haplotypes. Statistical significance of percent actuation was tested by Fisher’s exact test with Benjamini-Hochberg FDR correction at all FIRE peaks identified for the normal haplotype (Gen I expanded had no significant differences within this window). The bottom three panels represent Fiber-seq FIRE results for each group of haplotypes. ENCODE fibroblast DNase-seq and ChIP-seq tracks are shown below for reference.

**Figure S3.**
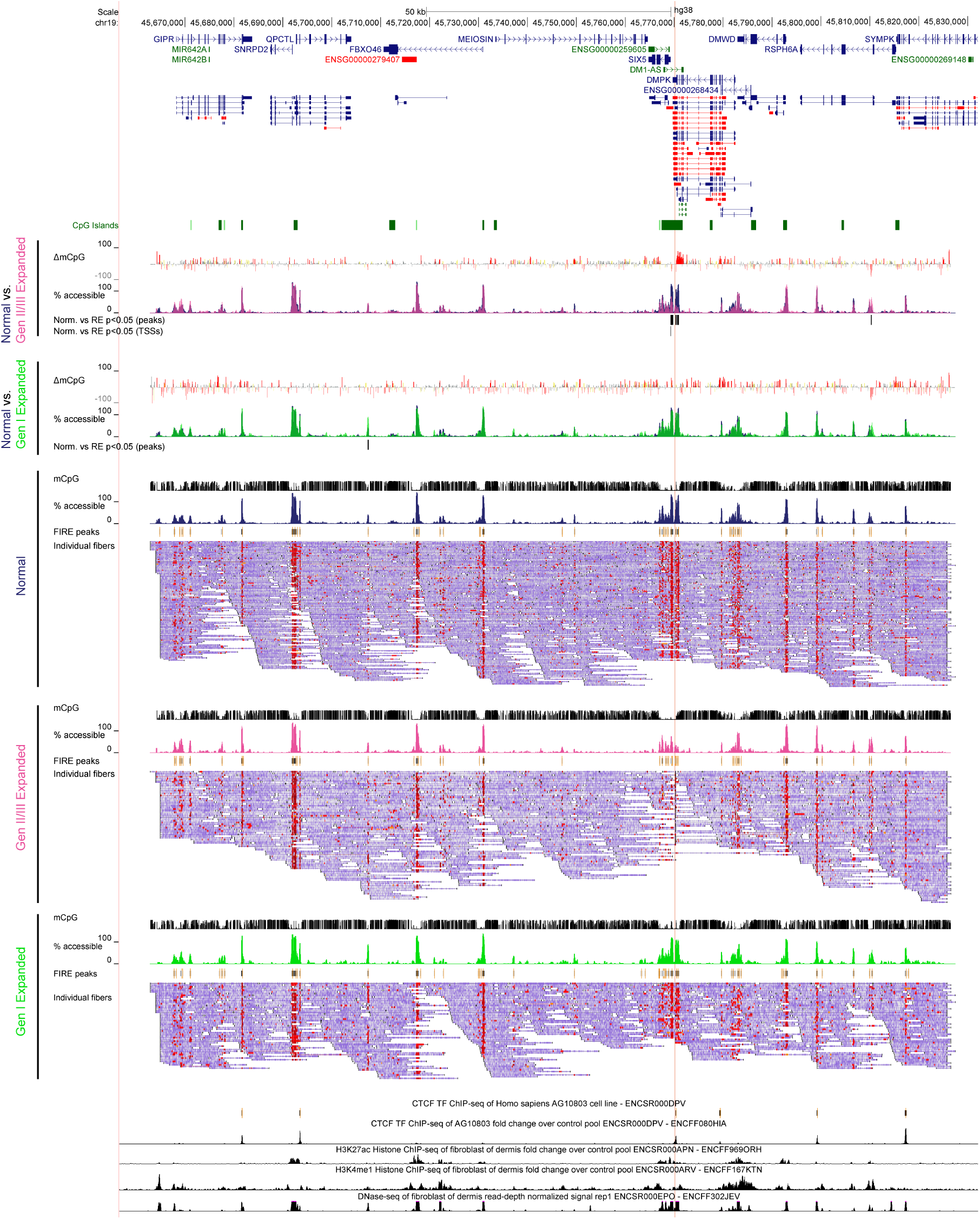
Targeted Fiber-seq across the targeted locus containing *DMPK*. A zoom-out of **Supplemental Fig. S2** across the entire targeted region. The “Normal vs. Gen II/III Expanded” panel also contains a track showing the *SIX5* promoter as the only GENCODE V46 annotated transcriptional start site with a statistically significant difference in percent actuation compared to normal haplotypes (Fisher’s exact test with Benjamini-Hochberg FDR correction). GENCODE V46 transcripts are shown above There were no significantly different TSSs for the Gen I expanded haplotype.

**Figure S4.**
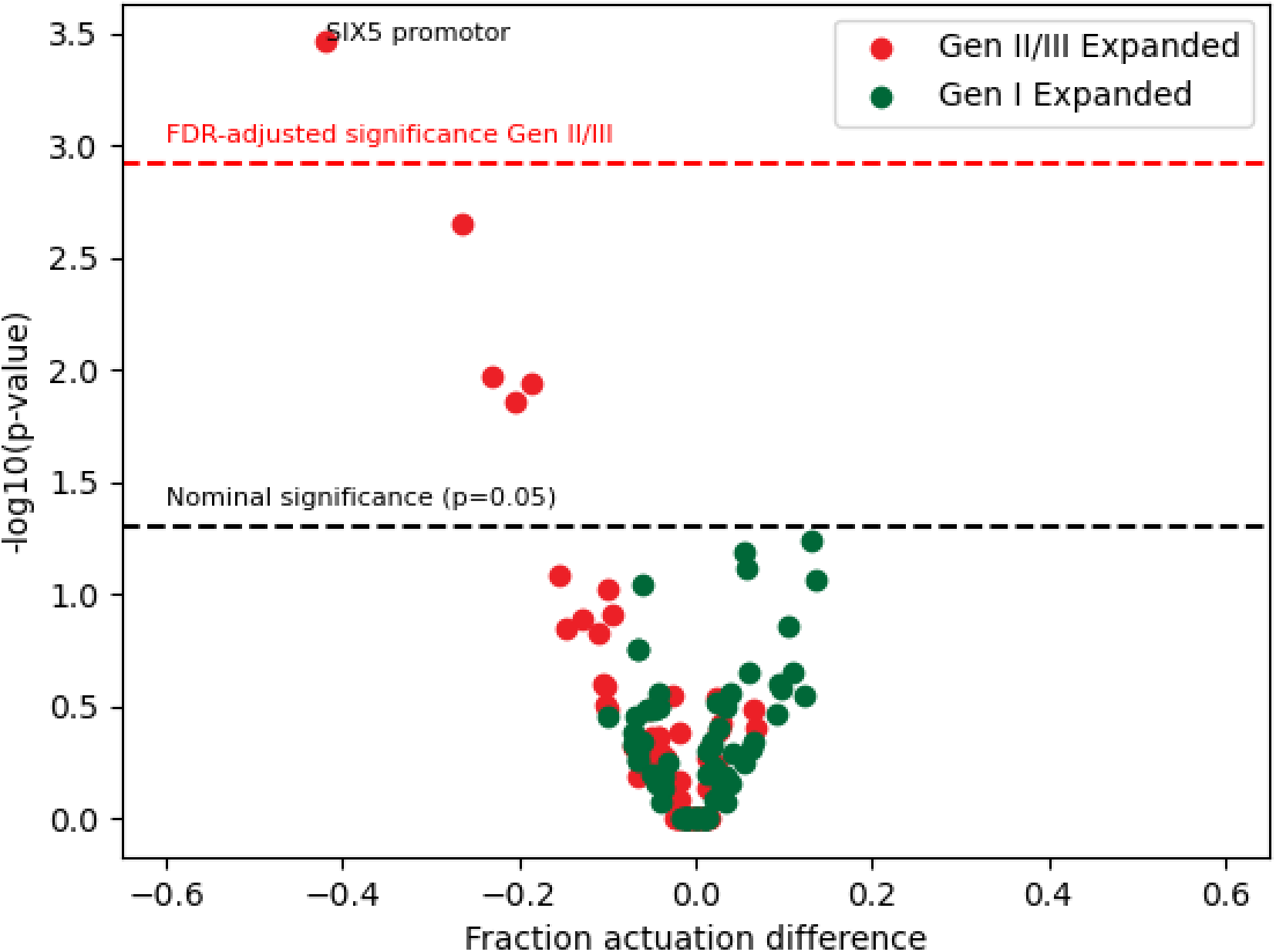
Volcano plot of actuation difference at all GENCODE V46 TSSs within the targeted locus. Y-axis p-values were calculated by Fisher’s exact test. The fraction actuation difference (x-axis) compares actuation among fibers from normal haplotypes to the Gen II/III expanded haplotypes (red) and Gen I expanded haplotype (green). The black dashed line denotes the threshold for statistical significance (p<0.05). Points above the red dashed line are significant with Benjamini-Hochberg FDR correction.

**Figure S5.**
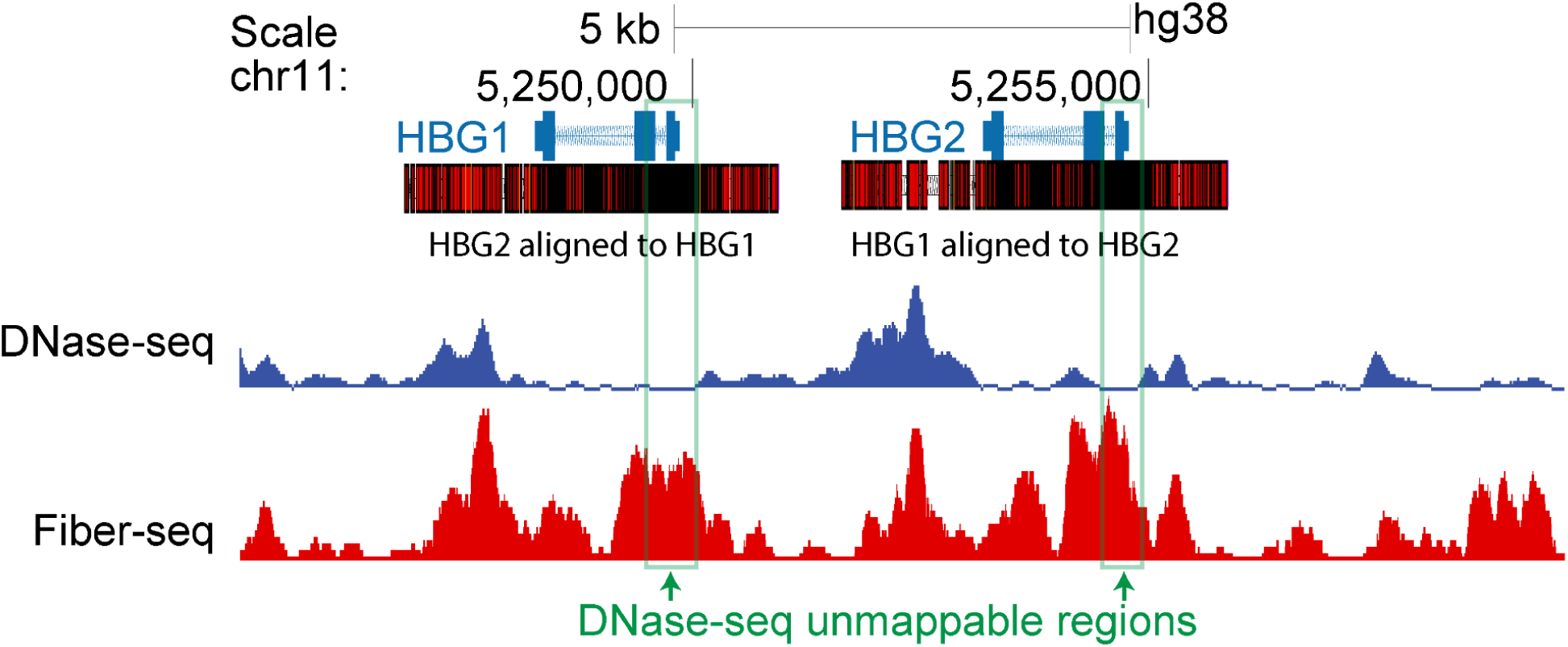
Sequence identity of *HBG1* and *HBG2*. Identical sequence is shown in black, with mismatches in red. Erythroid DNaseI-seq (blue, ENCODE track ENCFF800PMS) is shown above Fiber-seq percent actuated (red). Signal at the *HBG1*/*HBG2* promoters is missing from DNase-seq as short reads are not uniquely mappable due to 100% sequence identity.

**Figure S6.**
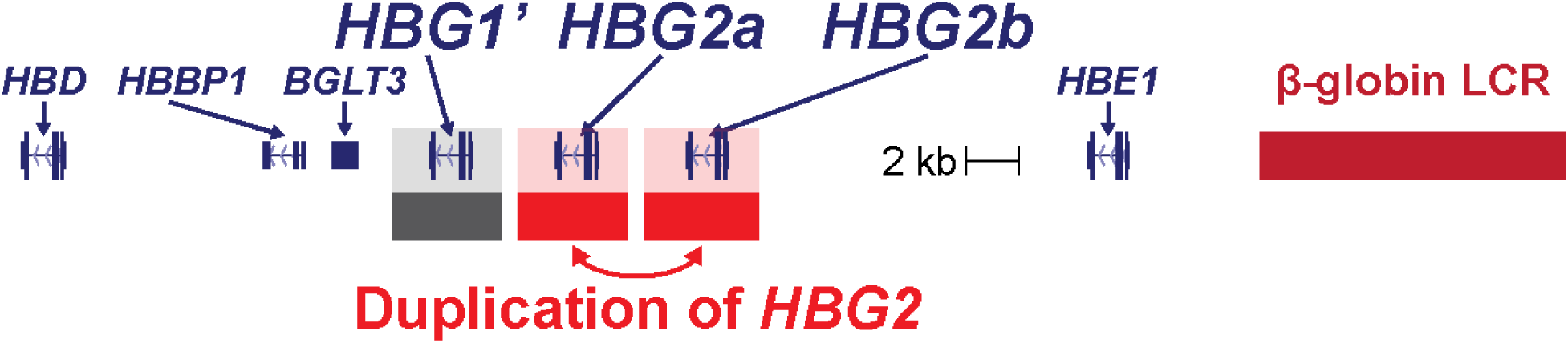
Schematic of the *HBG2* duplication in donor-derived CD34^+^ cells.

**Figure S7.**
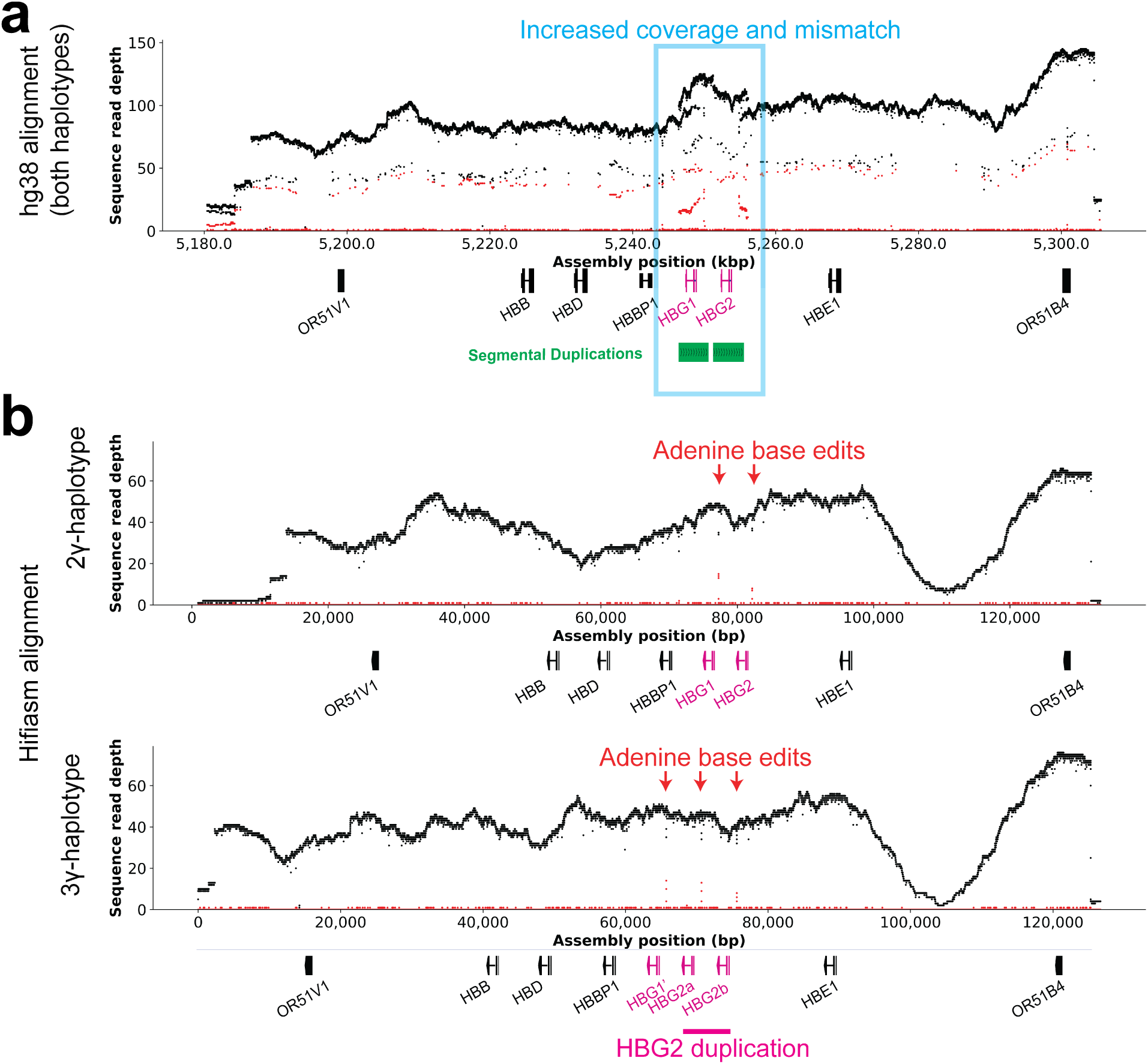
NucFreq plots (Vollger et al. 2019) of donor 1 at the β-globin locus. Read depth is shown in black, and mismatches are shown in red. **a)** Alignment to hg38, with the blue boxed region representing a spike in coverage and mismatch across the segmental duplications containing the *HBG1/HBG2* genes. **b)** Reads mapped to the diploid donor-specific assembly created with Hifiasm using HiFi reads from the same donor. Note that the spike in coverage and mismatch noted in a are resolved with inclusion of an apparent *HBG2* duplication event along the 3γ-haplotype.

**Figure S8.**
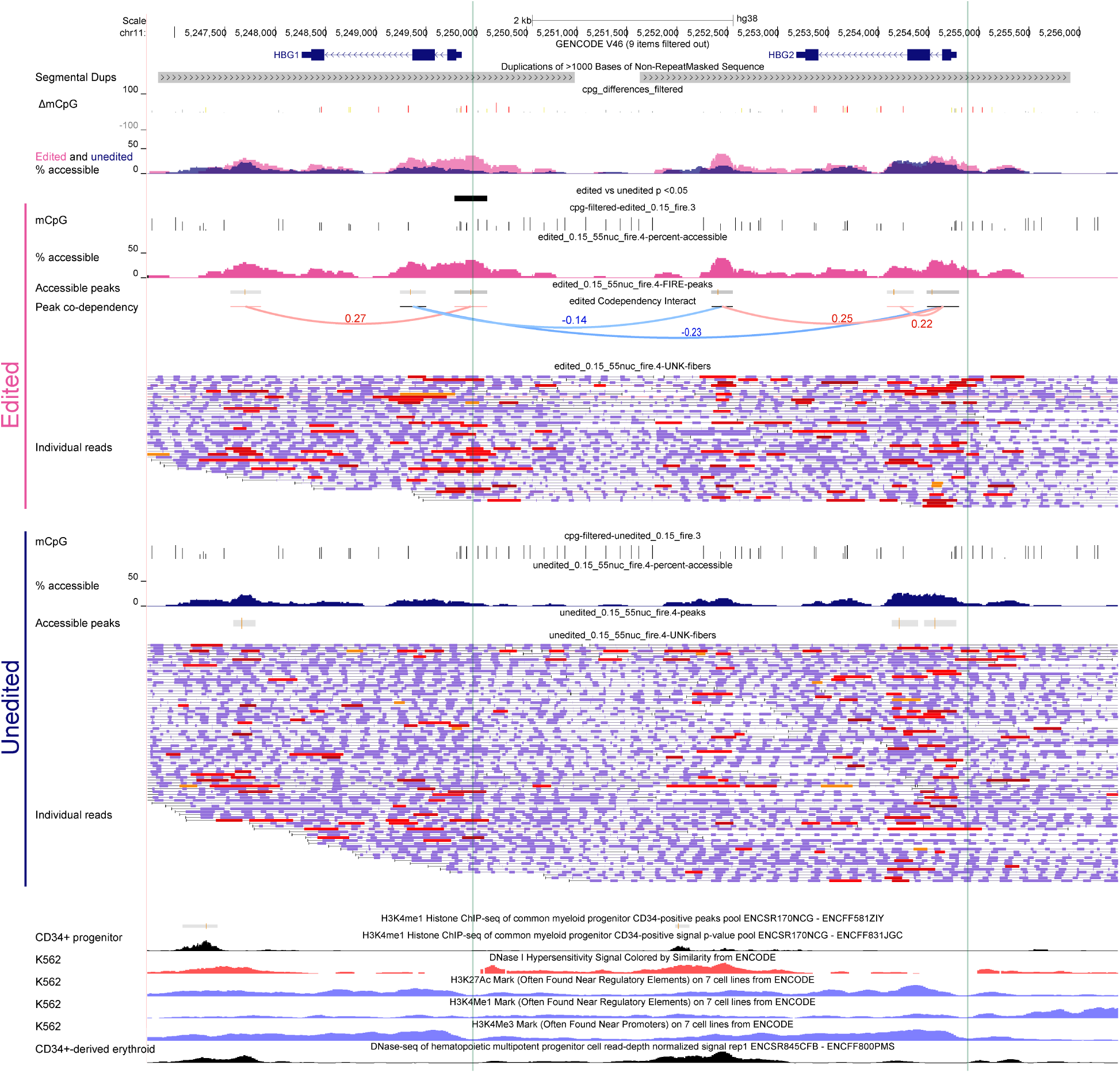
Targeted Fiber-seq across the γ-globin genes in ABE-edited CD34+ derived erythroid cells. Positive and negative codependencies of magnitude 0.1 or larger are displayed in red and blue, respectively. ABE-edited sites are highlighted in green. ENCODE tracks of H3K4me1 ChIP-seq from undifferentiated CD34^+^ progenitor cells, DNaseI and histone ChIP-seq from K562 cells (which express *HBG1*/*HBG2*), and DNaseI of CD34^+^ erythroid cells are included below.

**Figure S9.**
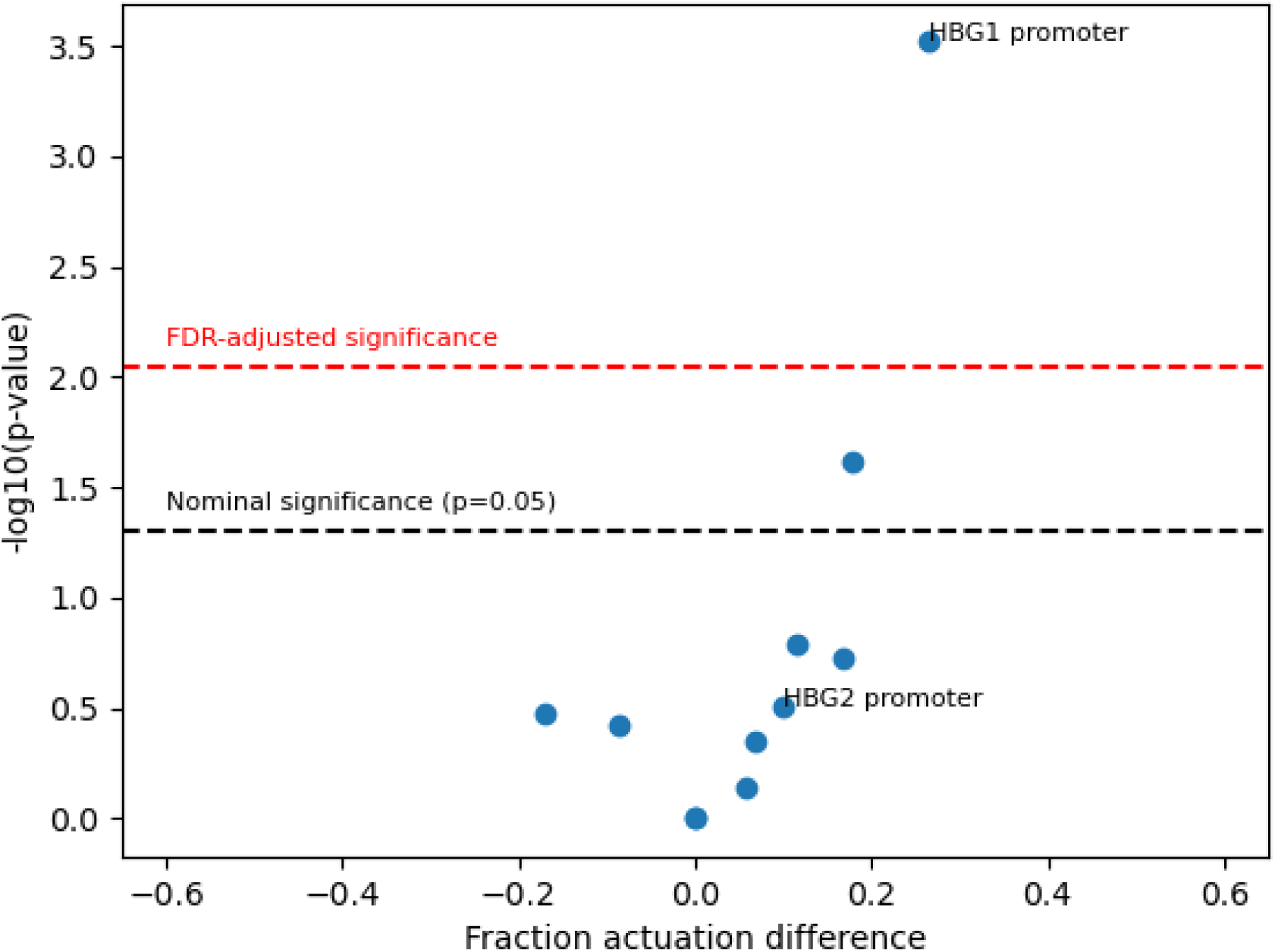
Volcano plot of actuation difference at all FIRE peaks from edited fibers. Y-axis p-values were calculated by Fisher’s exact test. The fraction actuation difference (x-axis) compares actuation among edited and unedited fibers from ABE-edited erythroid cells. The black dashed line denotes the threshold for statistical significance (p<0.05). Points above the red dashed line are significant with Benjamini-Hochberg FDR correction.

## Supplemental Tables

Supplemental Tables 1-2 can be found in the supplemental materials file called Supplemental_Tables.xlsx.

## Supplemental Data

Hifiasm assembly of the β-globin locus from the donor with the *HBG2* duplication is available on GitHub (https://github.com/StephanieBohaczuk/Targeted-Fiber-seq/tree/main/ABE_hifiasm_assembly).

## Supplemental Note

(1) The two peaks with increased actuation in expanded vs. normal haplotypes are both associated with variants that are consistent with the observed results:

Peak chr19:45,810,189-45,810,359 (significantly increased in Generations II/III)-There is a variant at chr19:45810295 (G/C). At this position, all normal haplotypes are C, and all expanded haplotypes from all generations are G (reference). This peak is also increased in Generation I but doesn’t meet significance.

Peak chr19:45707335-45707564 (significantly increased in Gen I): There is a variant at chr19:45707265 (C/T). At this position, all normal haplotypes are C (reference), the Generation I expanded haplotype is T, and the Gen II/III expanded haplotypes are C. This peak is not increased in the GenII/IIII expanded haplotype. Although the variant does not strictly overlap, the corresponding peak from the grandfather sample is wider (chr19:45,707,296-45,707,576) and there are several FIRE elements within the peak that extend to it.

## Notes

### Competing Interest Statement

Andrew Stergachis is a co-inventor on a patent relating to the Fiber-seq method (US17/995,058).
Andre Lieber is an academic co-founder of Ensoma Inc.
The remaining authors declare no competing financial interests.

